# Beyond category-supervision: instance-level contrastive learning models predict human visual system responses to objects

**DOI:** 10.1101/2021.05.28.446118

**Authors:** Talia Konkle, George A. Alvarez

## Abstract

Anterior regions of the ventral visual stream have substantial information about object categories, prompting theories that category-level forces are critical for shaping visual representation. The strong correspondence between category-supervised deep neural networks and ventral stream representation supports this view, but does not provide a viable learning model, as these deepnets rely upon millions of labeled examples. Here we present a fully self-supervised model which instead learns to represent individual images, where views of the same image are embedded nearby in a low-dimensional feature space, distinctly from other recently encountered views. We find category information implicitly emerges in the feature space, and critically that these models achieve parity with category-supervised models in predicting the hierarchical structure of brain responses across the human ventral visual stream. These results provide computational support for learning instance-level representation as a viable goal of the ventral stream, offering an alternative to the category-based framework that has been dominant in visual cognitive neuroscience.

## INTRODUCTION

Patterned light hitting the retina is transformed through a hierarchy of processing stages in the ventral visual stream, driving to a representational format that enables us to discriminate, identify, categorize, and remember thousands of different objects (Mishkin et al., 1983; Haxby et al., 2001; Kanwisher, 2010; DiCarlo and Cox, 2007; Grill-Spector and Weiner, 2014; Meyer and Rust, 2018). Prominent theoretical accounts of the organization of the high-level visual system assert that category-level (“domain-level”) forces are critical for shaping visual representation (Mahon and Caramazza, 2011; Peelen and Downing, 2017; Bracci et al., 2017; Op de Beeck and Ritchie, 2019; Kamps et al., 2020). Complementing this theoretical perspective, deep convolutional neural network models trained to perform multi-way object categorization learn hierarchical feature spaces that are currently the best predictive models of ventral visual stream responses to object images (Khaligh-Razavi and Kriegeskorte, 2014; Yamins et al., 2014; Güçlü and van Gerven, 2015; Cichy et al., 2016; Eickenberg et al., 2017; Wen et al., 2018; Schrimpf et al., 2018; Storrs et al., 2020, see Kriegeskorte, 2015; Serre, 2019 for review). However, it is clear that humans and non-human primates do not learn visual representation from millions of category labels, and that our perceptual systems discriminate visual objects without requiring category label information. These observations put into sharp focus a fundamental question about what the proximate goal of visual representation is — if not explicitly about object categories, what is an alternative, unsupervised, representational goal that would give rise to ventral stream representations?

One insight into this question, highlighted clearly by Wu et al. (2018), is based on the observation that the classification errors made by category-supervised networks are often semantically reasonable (e.g. an image of a leopard is more likely to be misclassified as a ‘jaguar’ than a ‘bookcase’). Critically, there is nothing explicit in the discriminative goal of multi-way object classification that enforces these relationships; instead these meaningful category-level relationships emerge through the natural covariance between visual features and broader conceptual divisions (c.f. Malcolm et al., 2016; Long et al., 2016, 2017). Wu et al. (2018) reasoned—and then demonstrated—that if the learning objective was changed to classify *each image*, then object category information would also implicitly emerge in the structure of the learned visual representation. Inspired by this instance-level supervised system, we developed a learning framework that is fully self-supervised, called instance prototype contrastive learning (IPCL), in which the goal is to learn a low-dimensional embedding of image-level representations. Further, because this instance-level learning framework can operate over any view of the world, regardless of what is depicted, it operationalizes a highly domain-general account of the pressures underlying the nature of visual tuning.

Here we consider instance-level contrastive learning as a proximate goal which can potentially learn generic representations that support downstream tasks such as object recognition and classification. Indeed, the field of machine learning uses object categorization capacity as a standard litmus test for whether a self-supervised model has learned generically useful representations (e.g. Chen et al., 2020a,b; Goyal et al., 2021), and this metric has also proven valuable in visual neuroscience research. In seminal work advocating for models with “performance-optimized” feature spaces, Yamins et al. (2014) highlighted a strong correspondence between a model’s ability to categorize objects and its ability to predict responses of individual neurons in object-selective inferotemporal (IT) cortex. This relationship between categorization capacity and brain predictivity has been more formally operationalized and expanded upon in the Brain-Score platform (Schrimpf et al., 2018). To date, considering a large number of category-supervised models, they find that gains in object categorization track strongly with gains in neural predictivity; however, recent models which are deeper and more accurate, no longer show increasingly brain-like representation (after about >70% top-1 accuracy on the ImageNet dataset; Deng et al., 2009). These performance-based relationships also raise a natural question for the present work—how strongly will models trained with instance-prototype contrastive learning show emergent object category structure, and how well will these features spaces show emergent brain-like representation, relative to their category-supervised counterparts?

Concurrently with the present work, and at a rapid pace, new models trained with different forms of instance-level learning have now become the state-of-the-art in self-supervised representation learning, with emergent object categorization capacity that rivals category-supervised models (Zhuang et al., 2019; Tian et al., 2019; He et al., 2019; Chen et al., 2020b,a; Caron et al., 2020). Although these models differ in terms of their architectural and algorithmic details, they share a common objective of encoding images as the same across views and different from other images. As such, these models also provide us with larger set of instance-level contrastive learning models to examine the relationship between emergent categorization accuracy and brain predictivity.

## RESULTS

### Instance-prototype contrastive learning

We designed an instance-prototype contrastive-learning algorithm (IPCL) to learn a representation of visual object information in a fully self-supervised manner, depicted in **Figure 1A**. The overarching goal is to learn a low-dimensional embedding of natural images, in which sampled views of the same image are nearby to each other in this space and also separable from the embeddings of all other images.

**Figure 1:**
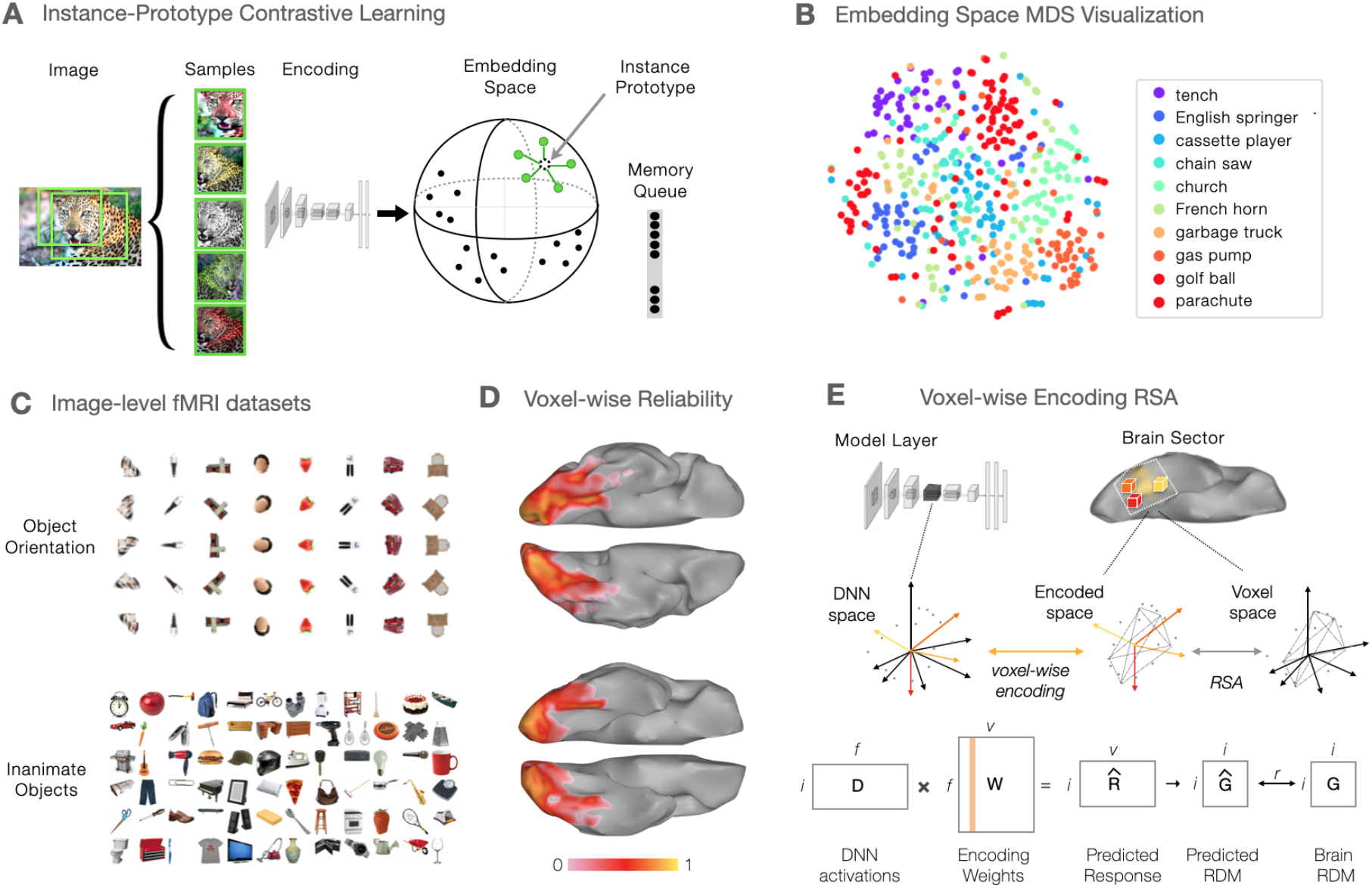
**A.** Overview of the self-supervised instance-prototype contrastive learning (IPCL) model which learns instance-level representations without category or instance labels. **B.** t-SNE visualization of 500 images from 10 ImageNet categories, showing emergent category clusters in deepnet feature space. **C.** All stimuli for the two fMRI datasets. Note that in this figure, the face image has been covered to remove identifying information. **D.** View from the bottom of the brain, showing voxel-wise reliability across the ventral visual stream for the Object Orientation dataset (top) and Inanimate Objects dataset (bottom). **E.** Overview of the voxel-wise encoding RSA procedure.

To do so, each image is sampled with 5 augmentations, allowing for crops, rescaling, and color jitter (following the same parameters as in Wu et al., 2018). These samples are passed through a deep convolutional neural network backbone, and projected into a 128-dimensional embedding space, which is L2-normed so that all image embeddings lie on the unit hypersphere. The contrastive learning objective has two component terms. First, the model tries to make the embeddings of these augmented views similar to each other by moving them towards the average representation among these views—the “instance prototype.” Simultaneously, the model tries to make these representations dissimilar from those of recently encountered items, which are stored in a light-weight memory queue of the most recent 4096 images—the “contrastive” component. See the **Supplementary Information** for the more precise mathemetical formulation of this loss function.

For the convolutional neural network backbone, we used an AlexNet architecture (Krizhevsky et al., 2012), modified to have group-normalization layers rather than standard batch normalization; see **Supplementary Figure 1**), which was important to stabilize the learning process. While traditional batch normalization operates by normalizing across images for each feature channel (Ioffe and Szegedy, 2015), group normalization operates by normalizing across groups of feature channels for each image (Wu and He, 2018), with intriguing parallels to divisive normalization operations in the visual system (Heeger, 1992; Carandini and Heeger, 2012). Five IPCL models were trained under this learning scheme, with slightly different training variations; all training details can be found in the **Supplementary Information**.

### Emergent Object Category Information

To examine whether these self-supervised models show any emergent object category similarity structure in the embedding space, we used two standard methods to assess 1000-way classification accuracy on ImageNet. The k-nearest neighbor (kNN) method assigns each image a label by finding the *k* (=200) nearest neighbors in the feature-space, assigning each of the 1000 possible labels a weight based on their prevalence amongst the neighbors (scaled by similarity to the target), and scoring classification as correct when the top-weighted class matched the correct class (top-1 knn accuracy; Wu et al., 2018). The linear evaluation protocol trains a new 1000-way classification layer over the features of the penultimate layer to estimate how often the top predicted label matches the actual label of each image (see Chen et al., 2020a,b; see **Supplementary Information** for method details).

Object category read-out from the primary IPCL models achieved an average top-1 kNN accuracy of 37.3% (35.4 – 38.4%) from the embedding space, and 37.1% (32.2 – 39.7%) from the penultimate layer (fc7). In contrast, untrained models with a matched architecture show minimal object categorization capacity, with top-1 kNN accuracy of 3.5% (3.3 – 3.8%) and top-1 linear evaluation accuracy of 7.2% (fc7). **Figure 1B** visualizes the category structure of an IPCL model, showing a t-SNE plot with a random selection of 500 images from 10 categories, arranged so that images with similar IPCL activations in the final output layer are nearby in the plot. It is clear that images from the same category cluster together. Thus, these fully self-supervised IPCL models have learned a feature space which implicitly captures some object category structure, with no explicit representational pressure to do so.

For comparison, we trained a category-supervised model with matched architecture and visual diet, and tested the categorization accuracy with the same metrics as the self-supervised model. The kNN top-1 accuracy was 58.8%, with a linear readout of 55.7% from the penultimate layer (fc7). An additional category-supervised matched-architecture model, trained with only one augmentation per image (rather than 5, which is a more standard training protocol), also showed similar classification accuracy (readout from fc7: kNN top-1: 55.5%; linear evaluation top-1: 54.5%). Thus, these matched-architecture category supervised models have noteably better categorization accuracy on the ImageNet database than our IPCL-trained models. **Supplementary Table 1** reports the categorization accuracies for all of the individual models.

### Relationship to the structure of human brain responses

To the extent that categorization capacity is indicative of brain-like representation in this accuracy regime (e.g. Schrimpf et al., 2018), we would expect these fully self-supervised models to have feature spaces with at least some emergent brain-like correspondence, but not as strong as category-supervised models. However, it is also possible that feature spaces learned in these self-supervised models have comparable or even more brain-like feature spaces than category-supervised models (e.g. if the instance-level representational goal more closely aligns with that driving visual system tuning). Thus, we next examined the degree to which the IPCL feature spaces have an emergent brain-like correspondence, relative to the category-supervised models.

Brain responses were measured using functional magnetic resonance imaging (fMRI) in two different condition-rich experiments (**Figure 1C**, see **Methods** and **Supporting Information**). The *Object Orientation* dataset included images of 8 items presented at 5 different in-plane orientations; this stimulus set probes for item-level orientation tolerance along the ventral visual hierarchy, while spanning the animate/inanimate domain. The *Inanimate Objects* dataset included images of 72 everyday objects; this stimulus set probes finer-grained distinctions within the inanimate domain. Thus, these two stimulus sets provide complementary views into object similarity structure. The resulting data revealed reliable voxel-level responses along the ventral visual stream (**Figure 1D**; see **Methods**). To delineate brain regions along the hierarchical axis of the ventral stream, we defined three brain sectors reflecting the early visual areas (V1-V3), the posterior occipito-temporal cortex (pOTC), and the anterior occipito-temporal cortex (aOTC; see **Methods**). Within these sectors, the group-averaged representational geometries were also highly reliable (EarlyV split-half reliability: r=.86-.90; pOTC: r=.75-.90; aOTC: r=.60-.89), providing a robust target to predict with different deep neural networks.

To relate the representations learned by these deep neural networks with brain sector responses along the ventral visual hierarchy, we used an approach that leveraged both voxel-wise encoding methods (Mitchell et al. 2008; Naselaris et al. 2011) and representational similarity (Kriegeskorte et al. 2008), which we subsequently refer to as voxelwise-encoding RSA (veRSA; **Figure 1E**; see **Methods**; see also Khaligh-Razavi et al., 2017; Kriegeskorte and Wei, 2021). This method fits an encoding model at each voxel independently, using weighted combinations of deepnet units (*W*) to predict the univariate response profile. Then, the set of voxel encoding models are used to predict multi-voxel pattern responses to new items (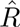) and to derive the predicted representational geometry in this encoded space (*Ĝ*). This predicted RDM is then compared to the RDM of the brain sector (*G*), as the key measure how well the layer’s features fit to that brain region. This analysis choice places theoretical value on the response magnitude of a voxel as an informative brain signature, while also reflecting the theoretical position in which neurons across the cortex participate as a unified population code.

The brain predictivity of the models are depicted in **Figure 2**. The results show that the IPCL model achieves parity with the category-supervised models in accounting for the structure of brain responses, evident across both datasets and at all three levels of hierarchy. Each plot shows the layer-wise correlations between the predicted and measured brain representational geometry, with all IPCL models in blue (with multiple lines reflecting replicates of the same model with slight training variations, see **Methods**), and category-supervised models in orange. The adjacent plots show the maximum model correlation, reflecting the layer with the strongest correlation with the brain RDM, computed with a cross-validated procedure to prevent double-dipping (cv max-r; see **Methods**), plotted for IPCL models, category-supervised models, and an untrained model. **Supplementary Table 2** reports the statistical tests comparing the brain predictivity between IPCL and category-supervised models, e.g. in 56/60 comparisons, the cross-validated max correlation for the IPCL models is greater than or not significantly different from category-supervised models (and with Bonferroni correction for multiple comparisons category-supervised models never showed a significantly higher correlation than an IPCL model).

**Figure 2:**
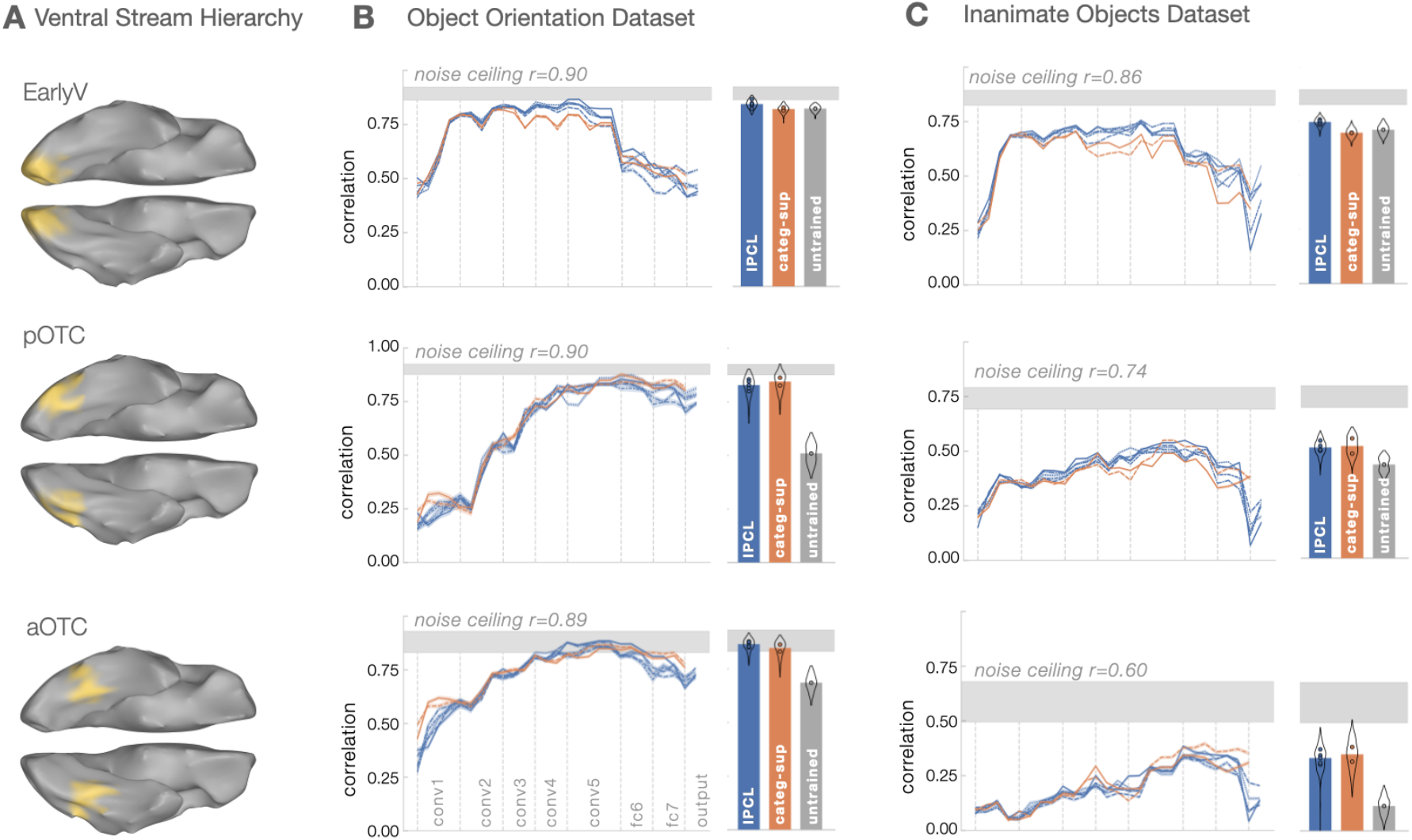
Model-to-brain fits. **A.** Visualization of the ventral stream regions of interest spanning the visual hierarchy from posterior to anterior (EarlyV, pOTC, aOTC). **B** and **C** show the veRSA results for the Object Orientation and Inanimate Object datasets, respectively. Each panel plots the correlation between model RDMs with neural RDMs (y-axis), computed separately for each model layer (x-axis) and brain region (rows). All IPCL models are in blue, and category-supervised models are in orange. The thickness of each line reflects 95% confidence intervals based on 1000 bootstrapped samples across split-halves. Bar plots show cross-validated estimates of the maximum correlation across model layers for each model class (IPCL in blue, category-supervised in orange, and an untrained model in gray). Error bars reflect a mirrored density plot (violin plot) showing the distribution of correlations across all split-halves, aggregated across instances of a given model type. Distributions are cutoff at ±1.5 IQR (interquartile range, Q3-Q1).

Further, all models account for a large proportion of the explainable variance in these highly-reliable brain representational geometries—though with a noticeable difference between the two datasets. Considering the *Object Orientation* dataset, the proportion of explainable variance accounted for approached the noise ceiling in all sectors for both IPCL and the category-supervised models (mean IPCL: 88%, 84%, 94%; category-supervised: 82%, 91%, 87%; noise ceiling: r=.90, .90, .89; for EarlyV, pOTC, and aOTC, respectively). However, considering the *Inanimate Objects* dataset, neither the IPCL nor category-supervised counterpart models learned feature spaces that reached as close to the noise ceiling, leaving increasing unaccounted for variance along the hierarchy (mean IPCL: 74%, 47%, 32%; category-supervised: 65%, 41%, 28%; noise ceiling: r=.86, .74, .60; for EarlyV, pOTC, aOTC, respectively). These results reveal that the particular stimulus distinctions emphasized in the dataset matter, as these dramatically impact the claim of whether the representations learned by these models are fully brain-like, or whether the models fall short of the noise ceiling.

Finally, these results also generally show a hierarchical convergence between brains and deep neural networks, with earlier layers capturing the structure best in early visual cortex, and later layers capturing the structure in the occipitotemporal cortex. However, unexpectedly, we also found that the untrained models were competitive with the trained models in accounting for responses in EarlyV and partially in pOTC, whereas both IPCL and category-supervised models clearly outperform untrained models in aOTC. Interestingly, the predicted representational distances in untrained models hover around zero, but nevertheless contain small differences that are consistent with the brain data. Further, the use of Group Normalization layers also boost untrained models—e.g. local Response Norm or Batch Normalization generally fit neural responses less well, particularly in early visual cortex (see **Supplementary Figure 2**). These findings highlight that there are useful architectural inductive biases present in untrained networks.

Overall, these results show that our instance-prototype contrastive learning models, trained without category-level labels, can capture the structure of human brain responses to objects along the visual hierarchy, on par with the category-supervised models. This pattern holds even in later stages of the ventral visual stream, where inductive biases alone are not sufficient to predict brain responses.

### Varying the visual diet

As some of the reliable brain responses in the later hierarchical stages of the *Inanimate Objects* dataset was unexplained, we next explored whether variations in the visual diet of the IPCL models might increase their brain predictivity. For example, the pressure to learn instance-level representations over a more diverse diet of visual input might result in richer feature representations that better capture the structure neural representations, particularly in the later brain stages reflecting finer-grained inanimate object distinctions. However, it is also possible that the relatively close-scale and centered views of objects present in the ImageNet database are critical for learning object-relevant feature spaces, and that introducing additional content (e.g. from faces and scenes) will detrimentally affect the capacity of the learned feature space to account for these object-focused brain datasets.

To probe the influence of visual diet, we trained 6 new IPCL models over different training image sets (**Figure 3A**; see **Methods**, **Supplementary Information**, **Supplementary Table 1**), and compared their brain-predictivity to the ImageNet trained baseline. First, because we made some changes to the image augmentations to accommodate all image sets, we trained a new baseline IPCL model on ImageNet. Second, we used object-focused images from a different dataset as a test of near-transfer (OpenImages; Krasin et al., 2017; Kuznetsova et al., 2020). The third dataset was scene images (Places2; Zhou et al. 2017) which we consider an intermediate-transfer test, as models trained to do scene categorization also learn object-selective features (Zhou et al., 2014). The fourth dataset was faces (VGGFace2; Cao et al. 2018), a far-transfer test that allows to explore whether a visual diet composed purely of close-up faces learns features that are sufficient to capture the structure of brain responses to isolated objects. The fifth dataset included a mixture of objects, faces, and places, which provides a richer diet that spans traditional visual domains, with the total number of images per epoch matched to the ImageNet dataset. The sixth dataset had the same mixture but used 3 times as many images per epoch to test whether increased exposure was necessary to learn useful representations with this more diverse dataset.

**Figure 3:**
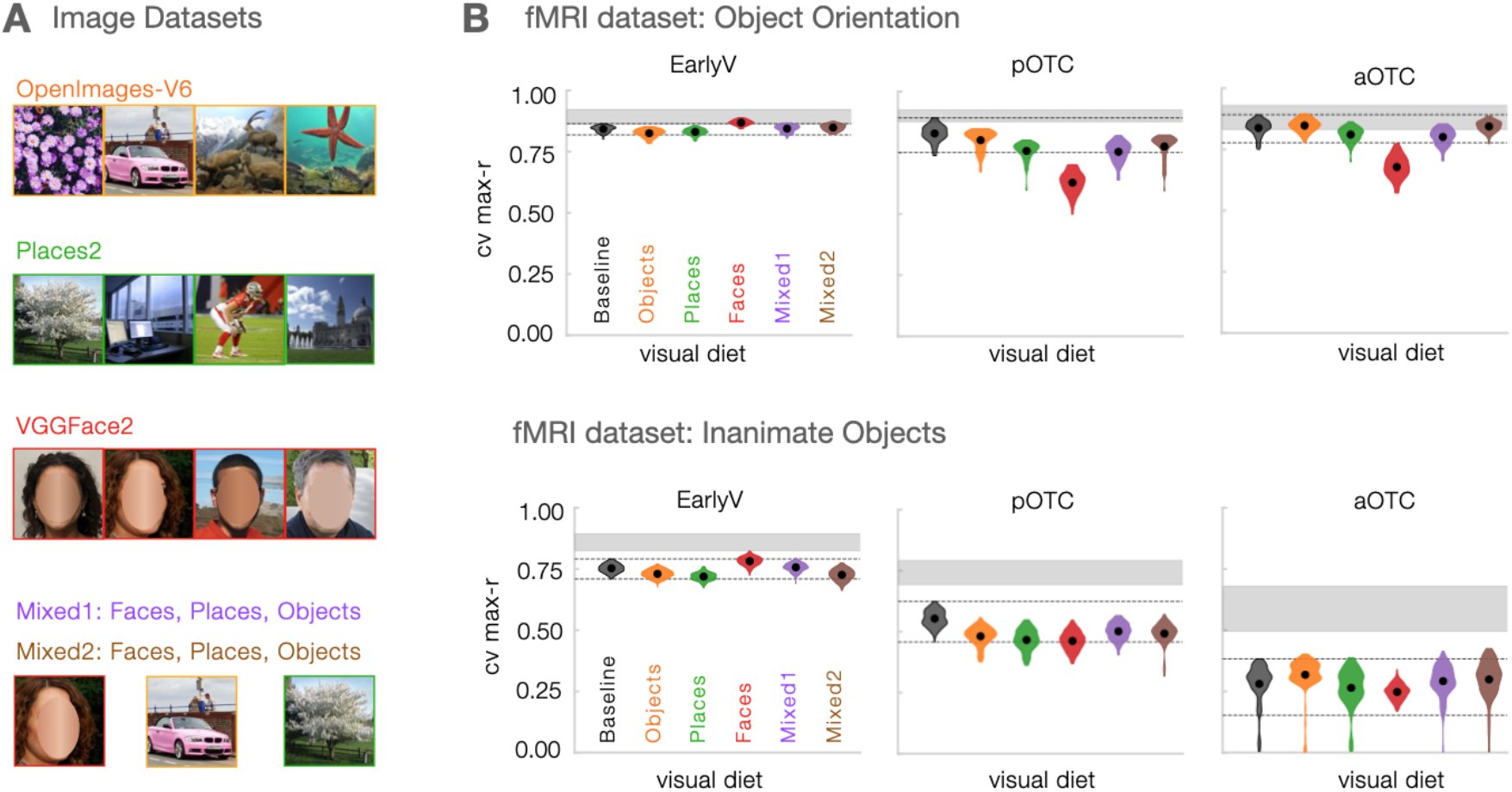
Consequences of variation in the visual diet. **A.** Example images are shown from each image dataset. Note that the faces shown are in the style of VGGFace2 (i.e., cropped views of faces), but are instead from thispersondoesnotexist.com, and are further covered to prevent identifying information. **B**. The cross validated maximum correlation (cv max-r) between model RDMs and neural RDMs for each dataset (rows), and each brain region (columns). Mean scores are shown with a black dot at the center of a mirrored density plot (violin plot) showing the distribution of correlations across all split-halves (distributions are cutoff at ±1.5 IQR, interquartile range, Q3-Q1). The dashed black lines indicate the ±1.5 IQR range for the matched baseline IPCL model trained on ImageNet.

For each of these six models trained with different kinds of visual experience, we used the same veRSA approach, and then calculated the cross-validated maximum correlation across layers (see **Methods**). The results are plotted in **Figure 3B**, where the five IPCL models with different visual experience (colored violin plots) are plotted in the context of the new baseline IPCL model trained on ImageNet (black dashed lines).

The overarching pattern of results shows that the visual diet actually had very little effect on how well the learned feature spaces could capture the object similarity structure measured in the brain responses. Quantitatively, the mean absolute difference in brain predictivity from the baseline ImageNet-trained model was Δ*r* <0.044 (range of signed differences −0.202 to 0.040). The visible outlier is the model trained only with views of faces. The features learned by this model were significantly less able to capture the structure of the *Object Orientation* dataset in both the posterior and anterior occipitotemporal cortex, with a difference from the baseline model >2.5 standard deviations from the mean difference across all comparisons (pOTC: z=3.67; aOTC: z=3.21). However, the feature spaces of this model were still able to capture the differences among objects in the *Inanimate Object* dataset, on par with the other visual diet variants in EarlyV and pOTC (though with a small reliable difference in pOTC) and was not different from the ImageNet trained baseline in aOTC (corrected t <1). The full set of results are reported in **Supplementary Table 3**.

Overall, this second set of IPCL models suggest that the statistics of most natural input contains the relevant relationships to comparably capture these brain signatures. Further, these models also highlight the general nature of the learning objective, demonstrating that it can be applied over richer and more variable image content, which is traditionally learned separately in supervised learning.

### Accuracy vs brain predictivity

The analyses so far demonstrate that, while category-supervised models show better object categorization capacity, IPCL models still achieve parity in their correspondence with the visual hierarchy. However, neither the category-supervised nor the IPCL models are able to fully capture the structure of the measured brain responses, particularly in the later hierarchical stage of the *Inanimate Objects* dataset that captures many finer-grained object relationships. This predictivity gap raises a new question — if instance-level contrastive learning systems advance to the point of achieving comparable emergent classification accuracy to category-supervised models, will even more brain-like representation emerge?

Concurrently, a number of new instance-level contrastive learning models have been developed, which allow us to test this possibility (e.g., SimCLR: Chen et al., 2020a; MoCo, MoCoV2: He et al., 2019; Chen et al., 2020b; SwAV: Caron et al., 2020). For example, SimCLR leverages related principles as our IPCL network, with a few notable differences: it uses two augmentations per image (rather than an instance prototype), a more compute intensive system for storing negative samples (in contrast to our light-weight memory queue), and a more powerful architectural backbone (Resnet50: (He et al., 2016)). Critically, this model, and others like MoCoV2 and SwAV, now achieve object classification performance that rivals their category-supervised comparands. Do these models show more brain-like representation, specifically in their responses to inanimate objects, where the later hierarchical brain structure was reliable and unaccounted for?

The results indicate that these newer models do not close this gap. **Figure 4** depicts the relationship between top-1 accuracy and the strength of the brain correspondence, for the *Inanimate Object* dataset. All instance-level contrastive learning models are plotted with colored markers, while category-supervised models are plotted with open markers. Different base architectures are indicated by the marker shape). These scatter plots highlight that, across these models, top-1 accuracy ranges from the 26-73%; however, improved categorization capacity is not accompanied by a more brain-like feature space. Further, these plots suggest that these particular variations in architecture, including higher powered ResNet (He et al., 2016) and ResNeXt (Xie et al., 2017) models, also do not seem to close this gap.

**Figure 4:**
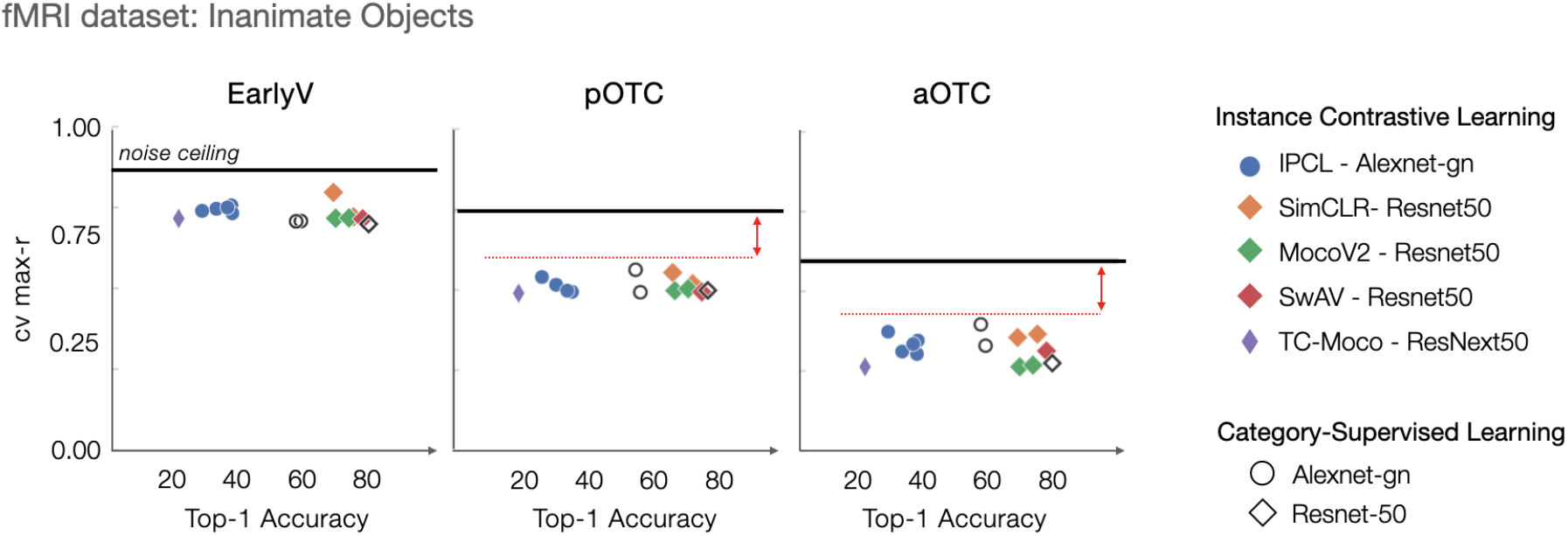
Relationship between object classification accuracy and brain correspondence. The x-axis plots top-1 classification accuracy, and the y-axis plots the cross-validated max correlation with the Inanimate Object dataset, in each of the three brain sectors. Self-supervised contrastive learning models are shown with colored markers, and category-supervised with open markers. Model architecture is indicated by marker shape. Red dashed line and double headed arrows draw attention to the gap between these model fits and the reliability ceiling of these brain data.

Finally, we also asked whether a recent self-supervised model trained on an even more ecological visual diet—images sampled from baby head-mounted cameras—might show better brain predictivity (TC-Moco: Orhan et al., 2020; SAYCam dataset: Sullivan et al., 2020). The visual experience of toddlers involves extensive experience with a very few things, rather than equal distribution over many categories–a visual curriculum which may be important for visual representation learning (Smith and Slone, 2017). However, this particular model also did not close the brain predictivity gap evident in the similarity structure of inanimate objects of at the later stages of the visual hierarchy (**Figure 4**; purple diamond). Note though that this model does not yet take advantage of temporal information in videos beyond a few frames; building effective systems that use contrastive learning over video is an active frontier (Sermanet et al., 2018; Zhuang et al., 2020; Knights et al., 2021).

Overall, the *Inanimate Objects* dataset has revealed some reliable representational structure in object-selective cortex that is not easily captured by current deepnet models, even across these broadly sampled variations in learning algorithm, architecture, and visual diet. Further, these aggregated results complement the emerging trend that overall object categorization accuracy is not indicative of overall brain predictivity (Schrimpf et al., 2018), here considering a variety of other instance-level contrastive learning methods, over a much wider range of top-1 accuracy levels.

### Auxiliary Results

For reference, we also conducted the same analyses using a classic representational similarity analysis (rather than veRSA), in which there was no voxel-wise encoding models, nor any deepnet unit feature re-weighting (**Supplementary Figures 2–4**). Overall, the magnitude of the correlation between model layers and the brain RDMs was systematically lower than when using veRSA. Despite this general main effect, the primary claims were also evident in this simpler analysis method: IPCL models showed parity with (or even superior performance to) category-supervised models, across brain sectors and datasets, with one notable exception. That is, in the aOTC and when considering the *Object Orientation* dataset, the category-supervised model showed better correspondence with the brain than the IPCL models (**Supplementary Figure 3**). This discrepancy between classic RSA and veRSA does highlight that veRSA is able effectively to adjust the representational space to better capture the brain data, while classic RSA weights all features equally. We discuss these results in the context of the open challenge of linking hypotheses between deepnet features and brain responses.

## DISCUSSION

Here we introduced instance-prototype contrastive learning models, trained with no labels of any kind, which learn a hierarchy of visual feature spaces that predict the representational geometry of hierarchical ventral visual stream processing in the human brain, on par with category-supervised counterparts. This result held in two datasets, considering both representational similarity among orientation variation, and with finer-grained inanimate object distinctions. By moving towards instance-level representation, this learning framework can operate over rich visual input without presupposing categories. And, we demonstrate this capacity by training IPCL on a variety of different visual diets, which continue to show emergent brain-like feature spaces even with increasing variety. Further, we find that in IPCL models, category-level similarity naturally emerges in the latent space, but also that increasingly accurate object categorization accuracy on ImageNet does not predict increasing brain-like representation in these datasets. Finally, we highlight that there is representational structure in the brain that was not well accounted for by any model tested, particularly in the anterior region of the ventral visual stream, related to finer-grained differences among inanimate objects. Broadly, these results provide computational plausibility for instance-level separability-—that is, to tell apart every view from every other view–as a plausible goal of ventral visual stream representation, which reflects a shift away from the category-based framework that has been dominate in high-level visual cognitive neuroscience research.

### Implications for the biological visual system

The primary advance of this work for insights into the visual system is to make a computationally supported learnability argument: it is possible to achieve some category-level similarity structure without presupposing explicit category-level pressures. Items with similar visual features are likely to be from similar categories, and we show that the goal of instance-level representation allows that natural covariance of the data to emerge in the latent space of the model — a result that is further supported by the expanding set of self-supervised models with emergent object categorization accuracy comparable to category-supervised systems (Tian et al., 2019; He et al., 2019; Chen et al., 2020b,a; Caron et al., 2020). Our work adds further support for the biological plausibility of the hypothesis by demonstrating an emergent correspondence with the similarity structure measured from brain responses—e.g. it is not the case that our self-supervised models learn a representation format that is decidedly un-brain-like. Indeed, recent work suggests that not all self-supervised learning objectives achieve brain-like representation with parity to category-supervised models (Zhuang et al., 2021).

Our model invites an interpretation of the visual system as a very domain-general learning function, which maps undifferentiated, unlabeled input into a useful representational format. On this view, the embedding space can be thought of as purely *perceptual interface*, with useful visual primitives over which separate conceptual representational systems can operate. For example, explicit object category level information may be the purview of more discrete compositional representational systems, that can provide ‘conceptual hooks’ into to different parts of the embedding space (c.f. Konkle et al., 2010; Gärdenfors, 2019). Intriguingly, new theoretical work suggests that instance-level contrastive learning may actually implicitly be learning to invert the generative process (i.e. mapping from pixels to the latent dimensions of the environment which give rise to the projected images; Zimmermann et al., 2021), suggesting that contrastive learning may be particularly well-suited for extracting meaningful representations from images.

What does the failure of these models to predict reliable variance in aOTC for the *Inanimate Objects* dataset tell us about the nature of representations in this region? Using this same brain dataset, we have found that behavioral judgments related to the shape similarity, rather than semantic similarity, show better correspondence with aOTC (Magri and Konkle, 2020, see also Baldassi et al., 2013; Jozwik et al., 2016). This result raises the possibility that the deepnets tested here are missing aspects of shape reflected in aOTC responses (e.g. structural representations: Lescroart and Biederman, 2013; global form: Ostwald et al., 2008; or configural representations: Wilson and Wilkinson, 2015), which resonates with the fact that CNNs operate more as local texture analyzers (Geirhos et al., 2018; Brendel and Bethge, 2019), and may be architecturally unable explicitly represent global shape (Doerig et al., 2020). Taken together, these results indicate that the success of CNNs in predicting ventral stream responses is driven by their ability to capture texture-based representations that are also extensively present throughout the ventral stream (Long et al., 2018), but they fall short where more explicit shape representations are emphasized. Capturing brain-like finer-grained distinctions among inanimate objects is thus as an important frontier that is currently beyond the scope of both contrastive and category-supervised models.

### Components of the learning objective

Why is instance-prototype contrastive learning so effective in forming useful visual representations, and what insights might this provide with respect to biological mechanisms of information processing? Recent theoretical work (Wang and Isola, 2020) has revealed that the two components of the contrastive objective function have two distinct and important representational consequences, which they refer to as alignment (similarity across views) and uniformity (using all parts of the feature space equally). To satisfy the alignment requirement, the model must learn what it means for images to be similar. For IPCL, the model takes 5 samples from the world, and tries to move them to a common part of the embedding space, forcing the model to learn that the perceptual features shared across these augmentations are important to preserve identity, while the unshared perceptual features can be discarded. Interpreted with a biological lens, these augmentations are like proto-eye movements, and this analogy highlights how this model can integrate more active sensing. For example, augmentations could sample over translation and rotation shifts of the kind that occur with eye and head movements. Further, “efference copy” signals (Colby et al, 1992; Crapse and Sommer 2007), which signal the magnitude and direction of movements between samples, might also lead to predictable shifts in the embedding space. This intrinsic information about the sampling process could enable the system to learn representations that are “equivariant”, as opposed to “invariant”, over identity-preserving transformations (c.f., Lenc and Vedaldi, 2015; Bouchacourt et al., 2021).

The second component of the objective function enforces representational uniformity–that is, where the set of all images have uniform coverage over the hypersphere embedding space. In IPCL this is accomplished by storing a modest set of “recent views” in a memory queue to serve as negative samples; other successful contrastive learning models use a much larger set of negatives (either in a batch or queue) which presumably helps enforce this goal (Chen et al., 2020a,b). The memory queue also has biological undertones: the human and non-human primate ventral streams are effectively a highway to the hippocampus (Van Essen and Maunsell, 1983). Through this lens, the recent memory queue of IPCL is a stand-in for the traces that would be accessible in a hippocampal memory system, inviting further modifications that vary the weight of the contrast with fading negative samples, or negative sample replay. However, we are not committed to memory queue data structure, *per se*. Given that its functional role is to give rise to good representational coverage over the latent space, there may be other architectural mechanisms by which the item separability can be achieved (Zbontar et al., 2021). Indeed, there is an ongoing debate about whether the instance-level separability requires these negative samples at all (Grill et al., 2020; Chen and He, 2020; Tsai et al., 2021).

While these instance-level contrastive learning systems advance a more biologically-plausible learning algorithm than category-supervised models, they are by no means a perfect model of how the brain learns–we instead see them as a testbed for broader learnability arguments and as useful for providing insights into visual representation and formats (e.g. clusters in an L2-normed hypersphere can easily be read-out with local linear hyperplanes, and this is not true of euclidean spaces; Wang and Isola, 2020), and as such serve as a useful computational abstraction.

### Concurrent work in non-human primate vision

In highly related recent work, Zhuang et al. (2021), explored a variety of self-supervised vision models and whether they have brain-like representation, using single-unit responses of the non-human primate ventral visual stream. Broadly, they found that the models using instance-level contrastive models achieved parity with category-supervised models in predicting responses in areas V1, V4, and IT; exceeding the capacities of other kinds of self-supervised models with different goals, including an autoencoder (a reconstructive goal), next frame prediction (PredNet: Lotter et al., 2020), and other non-relational objectives like depth labeling and colorization (Laina et al., 2016; Zhang et al., 2016). Further, they also capitalized on the value of this general objective, developing variations of their instance-level contrastive learning model to learn over video from the SAYcam baby head-cam dataset (Sullivan et al., 2020)–finding weaker but generally maintained neural predictivity. While almost every methodological detail is different from the work here, these two studies generally drive to very similar broad claims, arguing to move away from category-supervision towards instance-level contrastive learning. Further, the differences between our approaches reveal an expansive new empirical space to explore, considering different methods (fMRI, electrophysiology), models (IPCL, Local Aggregation), and model organisms (humans, monkeys); and, critically, the linking hypotheses (veRSA, encoding models) that operationalize our understanding of the neural code of object representation.

### Analytical Linking Hypothesis Between Model and Brain Activations

The question of how feature spaces learned in deep neural networks should be linked to brain responses measured with fMRI is an ongoing analytical frontier–different methods are abundant (e.g. Jozwik et al., 2017; Long et al., 2018; Eickenberg et al., 2017; Wen et al., 2018; Zeman et al., 2020; Storrs et al., 2020), each making different implicit assumptions about the nature of the link between model feature spaces and brain responses. In the present work, we assume a voxel is best understood as a weighted combination of deepnet features—-this is intuitive give the coarse sampling of a voxel over the neural population code. However, note that even single neuron responses (measured with electrophysiology in the primate brain) are modeled as weighted combinations of deepnet units, or even as weights on the principle components throughout the deepnet feature space (Klindt et al., 2017). In general, exactly how deepnet units are conceived of (e.g. how the tuning of any one deepnet unit is related to single neuron firing) is still coming into theoretical focus, where different hypotheses are implicit in the kind of regression model (e.g. whether encoding weights should be sparse and positive relationship, or low in magnitude and distributed across many deepnet units).

To arrive at a single aggregate measure of neural predictivity, encoding model approaches simply average across the set of individual neuron fits (e.g. Zhuang et al., 2021; Schrimpf et al., 2018). In contrast, we considered these voxel-wise encoding models together as an integrated population code, in which items vary in the similarity of their activation profiles, which focuses on the representational geometry of the embedding (Kriegeskorte et al., 2008). One motivation for this shift to the representational similarity as the critical neural target to predict is that fMRI allows for relatively extensive spatial coverage, providing access to a population-level code at a different scale than is possible with dozens to hundreds of single unit recordings; indeed trying to predict the RDM of a brain region is now the defacto standard in visual cognitive neuroscience. However, note that our approach differs from other kinds of weighted RSA analyses that are often employed on fMRI data (e.g. Jozwik et al., 2017; Storrs et al., 2020), which fit the representational geometry directly by re-weighting feature-based RDMs, discarding univariate activation profiles entirely. Finally, for RSA approaches, exactly how distances in a high-dimensional feature space are conceived of and computed is a further open frontier (Stringer et al., 2019), where different hypotheses about the way information is evident in the neural code are implicitly embedded in the choice of distance metrics (e.g. as the euclidean distance or the angle between vectors; e.g. Diedrichsen et al., 2020; Meyer and Rust, 2018).

At stake with these different analytical approaches is that the choices influence the pattern of results and subsequent inferences. For example, in the present data, model features are much more strongly related to brain RDMs when using veRSA than when using classic RSA, which make sense considering this method can recover true relationships that have been blurred by voxel-level sampling; however, untrained models also improve dramatically under this method, raising the question of whether the flexibility of re-weighting to the feature space is too great (or the Pearson-r scoring method is too lenient). As another example, in the present data, the IPCL features were able to comparably capture responses in aOTC to objects at different orientations, but only with veRSA, and not with classic RSA. This discrepancy between the analysis approaches suggests that the brain-like orientation information is embedded in the feature space, but requires voxel-wise encoding models to draw out those relationships—these pairwise relationships are less strongly evident in the unweighted feature space. Why? One possibility is that these IPCL models do not currently experience any orientation jitter across the samples (only crops, resizes, and coloration variation) and thus orientation-tolerance cannot enter into to the instance-prototype representations. In current work we are adding orientation augmentation to IPCL samples to explore this possibility. More broadly, we highlight these analytic complexities for two reasons. First, to be transparent about the untidy patterns in our data and the current state of our thinking for motivating these analysis decisions in the present work. And second, to open the conversation for the field to understand more deeply the ways in which deepnet models have brain-like representation of visual information under different analysis assumptions, especially as these new interdisciplinary analytical standard approaches are being developed.

### Conclusion

The prominence of category organization in the ventral visual stream has led to theories proposing that category-level (or “domain-level”) forces drive the organization of this cortex. That instance-level contrastive learning can result in emergent categorical representation supports an alternative theoretical viewpoint, in which category-specialized learning mechanisms are not necessary to learn representations with categorical structure. On this generalist account, visual mechanisms operate similarly over all kinds of input, and the goal is to learn hierarchical visual features that simply try to discriminate each view from every other view of the world, regardless of the visual content or domain. We further show that these instance-level contrastive learning systems can have representations that are as brain-like as category-supervised systems, increasing the plausibility of this general learning account. This generalist view does not deny the importance of abstract categories in higher level cognition, but instead introduces the instance-level learning objective as a proximate goal that learns compact representations that can support a wide variety of downstream tasks, including but not limited to object recognition and categorization.

## Methods

### Models

IPCL and category-supervised comparison models were implemented in PyTorch (Paszke et al., 2019), based on the codebase of Wu et al. (https://github.com/zhirongw/lemniscate.pytorch). Code and models available here: (https://github.com/harvard-visionlab/open_ipcl).

For our primary models, we trained 5 models with an Alexnet-gn architecture (**Supplementary Figure 1**), using instance-prototype contrastive learning (see **Supplementary Methods** for details), on the ImageNet-1k dataset (Deng et al., 2009). We used the data augmentation scheme used by Wu et al. (2018), with both spatial augmentation (random crop and resize; horizontal flip), and pixelwise augmentation (random grayscale; random brightness, contrast, saturation, and hue variation). These augmentations require the network to learn a representation that treats images as similar across these transformations. The replications reflect explorations through different training hyper-parameters. See the **Supplementary Methods** for extended details about the architecture, augmentations, loss function, and training parameters.

For the category-supervised model, we used the same AlexNet-gn architecture as in the primary IPCL models (minus the final L2-norm layer), but with a 1000-dimensional final fully-connected layer corresponding to the 1000 ImageNet classes. The standard cross-entropy loss function was used to train the model on the ImageNet classification task. Otherwise training was identical to the IPCL models, with the same visual diet (i.e., same batch size and number of augmented samples per image using the same augmentation scheme), and the same optimization and learning rate settings.

We trained 6 additional IPCL models to examine the impact of visual diet on learned representations, using datasets that focus on objects, places, faces, or a mixture of these image types: (1) ImageNet: ~1.28 million images spanning 1000 object categories (Deng et al., 2009). (2) Objects: OpenImagesV6, ~1.74 million training images spanning 600 boxable object classes (Krasin et al., 2017; Kuznetsova et al., 2020). (3) Faces: vggFace2, ~3.14 million training images spanning 8631 face identities (Cao et al., 2018). (4) Places: places2, ~1.80 million images of scenes/places spanning 365 categories; (Zhou et al., 2017), (5) Faces-Places-Objects-1x: a mixture of ImageNet, vggFace, and places2, randomly sampling images across all sets, limited to ~1.28 million images per epoch to match the size of the ImageNet training set, (6) Faces-Places-Objects-3x: limited to 3.6 million images per epoch. We used less extreme cropping parameters for all of these models than for the primary models so that the faces in the vggFace2 dataset would not be too zoomed in (as in this dataset, they tend to be already tightly cropped views of heads and faces). We used identical normalization statistics for each model (rather than tailoring the normalization statistics to each training set). Finally, we had to reduce the learning rate of the Faces model to .001 in order to stabilize learning. Otherwise, all other training details were identical to those for the primary models.

We also analyzed the representations of several concurrently-developed instance-level contrastive learning models: SimCLR: (Chen et al., 2020a), MoCoV2 (Chen et al., 2020b), and SwAV (Caron et al., 2020), which are trained on ImageNet; and TC-MoCo: (Orhan et al., 2020), trained on baby head-cam video data (Sullivan et al., 2020). These models were downloaded from official public releases.

To extract activations from a model, images were resized to 224 × 224 pixels and then normalized using the same normalization statistics used to train the model. The images were passed through the model, and activations from each model layer were retained for analysis. The activation maps from convolutional layers were flattened over both space and channel dimensions yielding a feature vector with length equal to NumChannels × Height × Width, while the output of the fully-connected layers provided a flattened feature vector with length equal to NumChannels.

### fMRI Experiments

The *Object Orientation* fMRI dataset reflects brain responses measured in 7 participants, while viewing images of 8 items presented at 5 different in-plane orientations (0, 45, 90, 135 and 180 degrees), yielding a total of 40 image conditions. These images were presented in a mini-blocked design, where in each 6min-12s run, each image was flashed 4 times (600ms on, 400ms off) in a 4s block, and was followed by 4s fixation. All 40 conditions were presented in each run; the order was determined using the optseq2 software, and was additionally constrained so that no item appeared in consecutive blocks (e.g. an upright dog, followed by an inverted dog). Two additional 20s rest periods were distributed throughout the run. Participants completed 12 runs. Their task was to pay attention to each image and complete a vigilance task (press a button when a red circle appeared around an object), which happened 12 times in run. Participants (ages 20-35, 4 female, unknown racial distribution) were recruited through the Department of Psychology at Harvard University, and gave informed consent according to procedures approved by the Harvard University Internal Review Board.

The *Inanimate Objects* fMRI dataset reflects brain responses measured in 10 participants, while viewing images depicted 72 inanimate items. In each 8-min run, each image was flashed 4 times (600ms on, 400ms off) in a 4s block, with all 72 images presented in a block in each run (randomly ordered), with 4 × 15s rest periods interleaved throughout. Participants completed 6 runs. Their task was to pay attention to each image and complete a vigilance task (press a button when a red-frame appeared around an object, which happened 12 times in run). Participants (ages 19-32; 8 females. unknown racial distribution) gave informed consent approved by the Institutional Review Board at the University of Trento, Italy.

Functional data were analyzed using Brain Voyager QX software and MATLAB, with standard preprocessing procedures and general linear modeling analyses to estimate voxel-wise responses to each condition at the single-subject level. Details related to acquisition and preprocessing steps can be found in the **Supplementary Information**.

#### Brain Sectors

First, the EarlyV sector was defined for each individual to include areas V1-V3, which were delineated based on activations from a separate retinotopy protocol. Next, an occipitotemporal cortex mask was drawn by hand on each hemisphere (excluding the EarlyV sector), within which the 1000-most active voxels were included, based on the contrast [all objects > rest] at the group-level. To divide this cortex into posterior and anterior OTC sectors, we used an anatomical cut off (TAL Y: -53), based on a systematic dip in local-regional reliability at a this anatomical location, based off of concurrent work also analyzing this *Inanimate Object* dataset (Magri and Konkle, 2020). The same posterior-anterior division was applied to define the sectors and extract data from the *Object Orientation* dataset.

#### Data reliability

The noise ceiling was defined in each sector, based on splitting participants into two groups, and averaging over all possible split halves. Specifically, we computed all of the subject-specific RDMs for each sector. Then, on a given iteration, we split the participants in half, and computed the average sector-level brain RDMs for each of these two groups. We computed the similarity of these two RDMs by correlating the elements along the lower triangular matrix (excluding the diagonal). The correlation distance (1-Pearson) was used for creating and comparing RDMs. This procedure was repeated for all possible split-halves over subjects. The noise ceiling was estimated as the mean correlation across splits (average of fisher-z transformed correlation values), and an adjusted 95% confidence interval that takes into account the non-independence of the samples (Bouckaert and Frank 2004). This particular method was used to dovetail with the model-brain correlations, described next.

### Model-Brain Analyses

The first key dependent measure (*veRSA correlation*) reflects the suitability of the features learned in a layer to predict the multivariate response structure in that brain sector. To compute this, we used the following procedure.

#### Voxelwise Encoding

For each deepnet layer, subject, and sector, each voxel’s response profile (over 40 or 72 image conditions, depending on the dataset) was fit with an encoding model. Specifically, in a leave-one-out procedure, a single image was held out, and ridge regression was used to find the optimal weights for predicting each voxel responses to the remaining images. We used sklearn’s (Pedregosa et al., 2011) cross-validated ridge regression to find the optimal lambda parameter. The response for the held-out item was then predicted using the learned regression weights. Each item was held out once, providing a cross-validated estimate of responses to each image in every voxel, which together form a model-based prediction of neural responses in each brain region. Based on these predicted responses, a model-predicted-RDMs was computed for each participant.

#### Layerwise RSA analysis

Next, for each sector and layer, the model-predicted-RDMs for each subject were divided into two groups and averaged, yielding two average model-predicted-RDMs from two independent halves of the data. Each RDM was correlated with actual brain-RDM, where the brain-RDM was computed from the same set of participants. This analysis was repeated for all possible splits-halves of the participants. The average fisher-z transformed correlation (and an adjusted 95% confidence interval Bouckaert and Frank 2004) was taken as the key measure of layer-sector correspondence.

Note that this average correlation reflects the similarity between the model-predicted-RDMs and the brain-RDMs, where only half of the subject’s brain data are used. This method of splitting the data into two halves was designed to increase the reliability in the data—we found that the RDMs were more stable with the benefit of averaging across subjects, while any one individual’s brain data was generally less reliable. Additionally, this procedure allows there to be some generality across subjects. Finally, we did not adjust the fit values to correct for the fact that the model-to-brain fit reflects only half the brain data, instead we kept it as is, which also allows the average layer-sector correlation to be directly compared to the similarly-estimated noise ceiling of the brain data.

#### Cross-Validated Max-Layer Estimation

The second key dependent measure relating model-brain correspondence reflects the strength of the best-fitting layer to a given sector. To compute this measure, we again used the same technique of splitting the data in half by two groups of subjects (this time to prevent double-dipping). Specifically, for each model and sector, the veRSA correlation was computed for all layers, and the layer with the highest veRSA correlation was selected. Then, in the independent half of the data (from new participants), the veRSA correlation was computed for this selected layer, and taken as a measure of the highest correspondence between the model and the sector. As above, this procedure was repeated for all possible split-halves of the subjects, and the cross-validated max-r measure was taken as the average across splits (averaging fisher-z transformed correlation values, and using the adjusted 95% confidence interval that takes into account the non-independence of the samples). This procedure insures an independent estimate of the maximum correspondence across layers.

#### Classic RSA

For comparison, we also computed and compared RDMs in both layerwise feature spaces and brain sectors using classic RSA. In this case, RDMs were computed directly from the deepnet activations (across units) and the brain activation patterns (across voxels), with no encoding model or feature weighting.

### Statistical Comparisons

To compare the cross-validated max correlation values between models, we used paired t-tests over all split halves of the data, with a correction for non-independence of the samples, following Bouckaert and Frank, 2004 (tests based on repeated k-fold cross validation) for corrected variance estimate and adjusted t-values. Comparisons between IPCL and Category-Supervised models are found in **Supplementary Table 2**; Comparisons between IPCL and an untrained model are found in **Supplementary Table 2**; Comparison between models trained with different visual diets to the baseline IPCL model trained on ImageNet are reported in **Supplementary Table 3**). Statistical significance for these paired t-tests was determined using a Bonferonni corrected *α* level of .05/30=0.00167, where 30 corresponds to the number of family-wise tests for all reported tests.

## Acknowledgments

Funding for this project was provided by an Amazon Cloud Credits for Research Grant to TK and GAA, an NSF CAREER BCS-1942438 to TK, and an NSF PAC COMP-COG 1946308 to GAA. Thank you to Morgan Henry who collected the Object Orientation dataset.

## Author Contributions

Both authors contributed extensively to this work. TK collected and pre-processed the Inanimate Object Dataset. TK and GAA supervised the collection and preprocessing of the Object Orientation dataset. TK organized all brain data for analysis. GAA implemented and trained all models. TK and GA jointly developed the self-supervised model, designed the experiments and analytical procedures, created the figures, and wrote the manuscript. The authors declare no competing interests.

## Supplementary Information

### 1 Extended Modeling Methods

#### 1.1 Instance Prototype Contrastive Learning

In contrastive-learning frameworks, the goal is to learn an embedding function that maps images into a low-dimensional latent space, where visually similar images are close to each other, and visually dissimilar images are far apart. Learning proceeds by organizing the training data into similar pairs (positive samples) and dissimilar pairs (negative samples), where different frameworks make different choices of how positive and negative samples are encoded and retained throughout the learning process.

In our instance-prototype contrastive learning framework, we randomly augment the same image (*x*) multiple times (*n* = 5 in the models reported here), then pass each augmented image (*x_i_* … *x_j_*) through an embedding function *f_θ_*(*x*) to obtain a low-dimensional representation of each image (*z_i_* … *z_j_*). We then compute an instance prototype 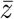 by averaging the embedding for all 5 samples:

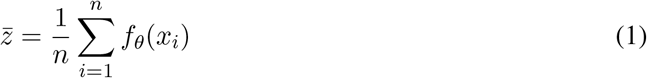

where n is the number of samples, *f_θ_*(*x*) is the embedding function (e.g., Alexnet-gn), and *x_i_* is the *ith* augmented sample of an image.

For each augmented instance, the prototype serves as its positive pair, and all stored representations serve as negative pairs (implemented with a lightweight, non-indexed memory queue storing the *K* = 4096 most recent samples). The normalized temperature-scaled cross entropy loss for a positive pair (*z_i_*, 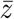) would be defined as:

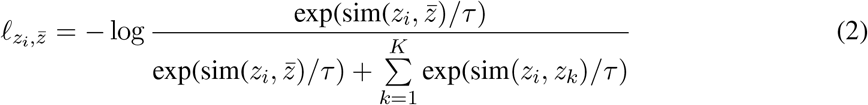

where the similarity function *sim* is the dot product between *L*2-normalized embeddings, *τ* is a temperature parameter that controls the dynamic range of the similarity function, and *K* is the total number of samples stored in the memory queue.

In practice, we used Noise Contrastive Estimation (NCE, Gutmann and Hyvärinen, 2010) to approximate sampling from a larger memory store (see Wu et al., 2018) though recent work suggests the loss function in equation 2 may suffice (Chen et al., 2020a). Specifically, we used Wu et al. (2018)’s implementation of Noise Contrastive Estimation to approximate sampling, with slight modifications to accommodate our prototype and queue:

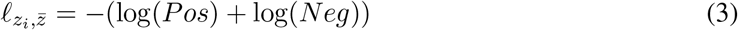

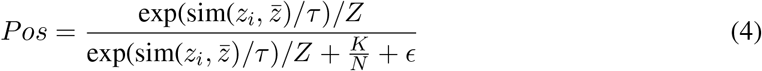

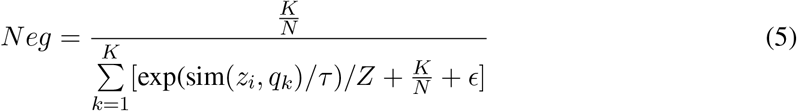

where *z_i_* is the embedding for the *i^th^* sample, 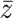 is its corresponding prototype, *sim* is the similarity function (dot-product between embeddings), *τ* is the temperature parameter, Z is a normalization constant (estimated based on the first mini-batch of 128 × 5 augmented samples), *q_k_* is the embedding for the *k^th^* item stored in the queue, and *ϵ* = 1e − 7 is a constant added for numerical stability.

The final loss is computed across all positive pairs in a minibatch (128 images, 5 samples per image, yielding 640 positive pairs). The queue is updated after every minibatch with the current samples added to the queue, displacing the oldest samples.

A PyTorch implementation and pretrained models for IPCL can be found at https://github.com/harvard-visionlab/open_ipcl.

#### 1.2 Model Architecture Details

We created a modified AlexNet-gn architecture following the original AlexNet implementation from Krizhevsky et al. (2012) with three noteworthy differences: (1) We used *group normalization* (Wu and He, 2018) with 32 groups per layer, instead of local response normalization with 5 channels per group (Krizhevsky et al., 2012). (2) For the self-supervised models, the final 1000-way output layer was replaced with a fully-connected low-dimensional embedding space (128 dimensions), followed by an L2-normalization layer necessary for the contrastive learning task. (3) The original AlexNet’s conv2, conv4, and conv5 layers were split across 2 GPUs for practical reasons — at the time GPUs had less RAM and could not fit the full model. While this split architecture can be emulated in PyTorch using grouped convolutional layers, we did not split the model architecture in this way because modern GPUs can fit the full model. Note the architecture details of this Alexnet are different from the official PyTorch version of AlexNet (https://pytorch.org/vision/stable/models.html), which implements Krizhevsky (2014). **Supplementary Figure 1** shows the exact model architecture specification for Alexnet-gn.

#### 1.3 Image Augmentation Details

Our primary models were trained with the following augmentations (“Aug Set1”): (1) RandomResizedCrop, which grabs a random crop from the original image with the scale restricted to (0.2,1.) times the area of the original image and an aspect ratio in the range (3/4,4/3) times the aspect ratio of the original image, and then this cropped image was resized to 224×224 pixels. (2) HorizontalFlip with probability=.5, (3) conversion to GrayScale with probability=.2, (4) RandomColorJitter which adjusted the brightness, contrast, and saturation between (.6,1.4) times the original, and hue between +/− 144 degrees of the original image. As is standard, images were also normalized by z-scoring each pixel (i.e., subtracting the mean and dividing by the standard deviation for each channel). Unless otherwise noted, we used the standard Imagenet normalization parameters (mean=[0.485, 0.456, 0.406], std= [0.229, 0.224, 0.225]).

For models varying the visual diet we used a slightly different augmentation setting (“Aug Set2”). Specifically, we reduced the RandomResizedCrop scale range to (0.5,1.0) because the face images were already relatively zoomed in. We also standardized the normalization parameters (mean=[0.5, 0.5, 0.5], std= [0.2, 0.2, 0.2]) to apply the same normalization to each dataset (ImageNet, OpenImagesV6, Places2, and VGGFace2).

Our models were trained with custom data-augmentation functions that operate on the GPU to accelerate augmentation.

#### 1.4 Training Details

Models were trained for 100 epochs with a batch size of 128×5 (128 images each augmented 5 times) using stochastic gradient descent, with momentum=.9 and weight decay=5e-4. Gradients were accumulated across 20 batches before each optimizer step. The learning rate was varied using the one-cycle policy (Smith, 2017), beginning at 0.00003, increasing with a cosine annealing function to a maximum of .03 after 40 epochs, then decreasing with a cosine annealing function toward zero (3e-09) by 100 epochs. As part of a hyperparameter search, three models were terminated after fewer epochs as their learning curves did not diverge from those trained with the parameters above: Alexnet-gn-ranger-ep82, trained with the Ranger optimizer (RAdam with Lookahead, Zhang et al. (2019)); Alexnet-gn-redux-73, trained with momentum on the same cosine annealing function as the learning rate, starting at .95, dropping to .85 after 40 epochs, then rising to .95); Alexnet-gn-transforms-82, trained with transforms customized to accelerate augmentation. Although these models weren’t trained for a full 100 epochs, they achieved similar top1 accuracy to models that were, and were therefore included in the primary analyses.

#### 1.5 Assessing Emergent Categorization Accuracy

##### 1.5.1 K-nearest neighbors evaluation

To classify a test image *x*, its embedding (e.g., 128 dimensional output activations) was compared to the embedding of each of the 1.28 million ImageNet training images using cosine similarity. The top *k* = 200 nearest neighbors were used to make the prediction via cosine-similarity-weighted voting, where the class c would receive the total weight given by:

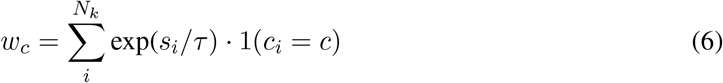

where *N_k_* denotes the k-nearest neighbors, and *s_i_* is the cosine similarity between the target and the neighbor, *k* = 200, and *τ* = 0.07 (matching the value used when computing the contrastive loss during training). The target was assigned to the class with the highest weight, and scored as correct/incorrect by comparing to the ground-truth label.

##### 1.5.2 Linear evaluation

To test whether category can be linearly decoded from model activations we trained a single fully connected layer with 1000 units on Imagenet-1k classification using the standard linear evaluation protocol (e.g., Chen et al. 2020b), in which we train a classifier on the output activations of the penultimate model layer (fc7 for Alexnet-gn models, the average pooling layer for resnet models). All parameters of the model being evaluated, including any normalization parameters, were frozen, and only the weights and biases of the fully-connected readout layer were updated. The standard linear evaluation protocol is slow and costly, training on ImageNet for 100 epochs, using stochastic gradient descent (momentum= 0.9, weight decay=0) and an initial learning rate of 30.0 which is reduced to 3.0 on epoch 60, then to .30 on epoch 80. To reduce training time we modified this standard linear evaluation protocol which enabled us to obtain similar performance levels in 10 epochs. Specifically, the learning rate was varied using the one-cycle policy (Smith, 2017), beginning at 0.00003, increasing with a cosine annealing function to a maximum of .3 after 3 epochs, then decreasing with a cosine annealing function toward zero (3e-09) by 10 epochs. We found top1 accuracy was often better, and certainly comparable with this more econimical procedure (e.g., standard vs. one-cycle top1 accuracy, 35.8% vs. 37.1% for IPCL Alexnet-gn readout from avgpool layer; 53.0% vs. 55.7% for category-supervised Alexnet-gn readout from fc7; 70.4% vs 72.0% for SWaV Resnet50 readout from the avgpool layer).

### 2 Extended fMRI methods

#### 2.1 Object Orientation Dataset

##### 2.1.1 MRI Acquisition

Imaging data were collected on a 3T Siemens Trio scanner at the Harvard University Center for Brain Sciences. Structural data were obtained in 176 axial slices with 1 × 1 × 1 mm voxel resolution, TR = 2200 ms. Functional blood oxygenation level-dependent (BOLD) data were obtained using a gradient-echo echo-planar pulse sequence (33 axial slices parallel to the anterior commissure-posterior commissure line; 70 × 70 matrix; FoV = 256 × 256 mm; 3.1 × 3.1 × 3.1 mm voxel resolution; gap thickness = 0.62 mm; TR = 2000 ms; TE = 60 ms; flip angle = 90 degrees). Volumes were acquired in ascending order. A 32-channel phased-array head coil was used.

##### 2.1.2 Data Pre-Processing

All fMRI data was processed using Brain Voyager QX software. Preprocessing steps included 3D motion correction, slice scan-time correction, linear trend removal, temporal high-pass filtering (0.01 Hz cutoff), spatial smoothing (4mm FWHM Kernel), and transformation into Talairach space. Statistical analyses were based on the general linear model. All GLM analyses included box-car regressors for each stimulus block convolved with a gamma-function to approximate the idealized hemodynamic response. All subsequent brain-based analyses were performed using the estimated beta coefficients from the single-subject voxel-wise GLMs.

#### 2.2 Inanimate Objects Dataset

##### 2.2.1 MRI Acquisition

Imaging data were acquired on a BioSpin MedSpec 4T scanner (Bruker) at the University of Trentro, Italy. Functional data were collected using an echo-planar 2D imaging sequence (TR, 2000ms; TE, 33ms; flip angle, 73°; slice thickness, 3mm; gap, 0.99mm, with 3 × 3 in-plane resolution). Volumes were acquired in the axial plane parallel to the anteroposterior commissure in 34 slices, with ascending interleaved slice acquisition.

##### 2.2.2 Data Pre-Processing

Functional data were analyzed using Brain Voyager QX software. Preprocessing included slice scan-time correction, 3D motion correction, linear trend removal, temporal high-pass filtering (0.01 Hz cutoff), spatial smoothing (6mm FWHM kernel), and transformation into Talairach (TAL) coordinates. General linear model analyses included square-wave regressors for each condition’s presentation times, convolved with a gamma function to approximate the hemodynamic response. All subsequent brain-based analyses were performed using the estimated beta coefficients from the single-subject voxel-wise GLMs.

### Supplementary Figures

**Supplementary Figure 1:**
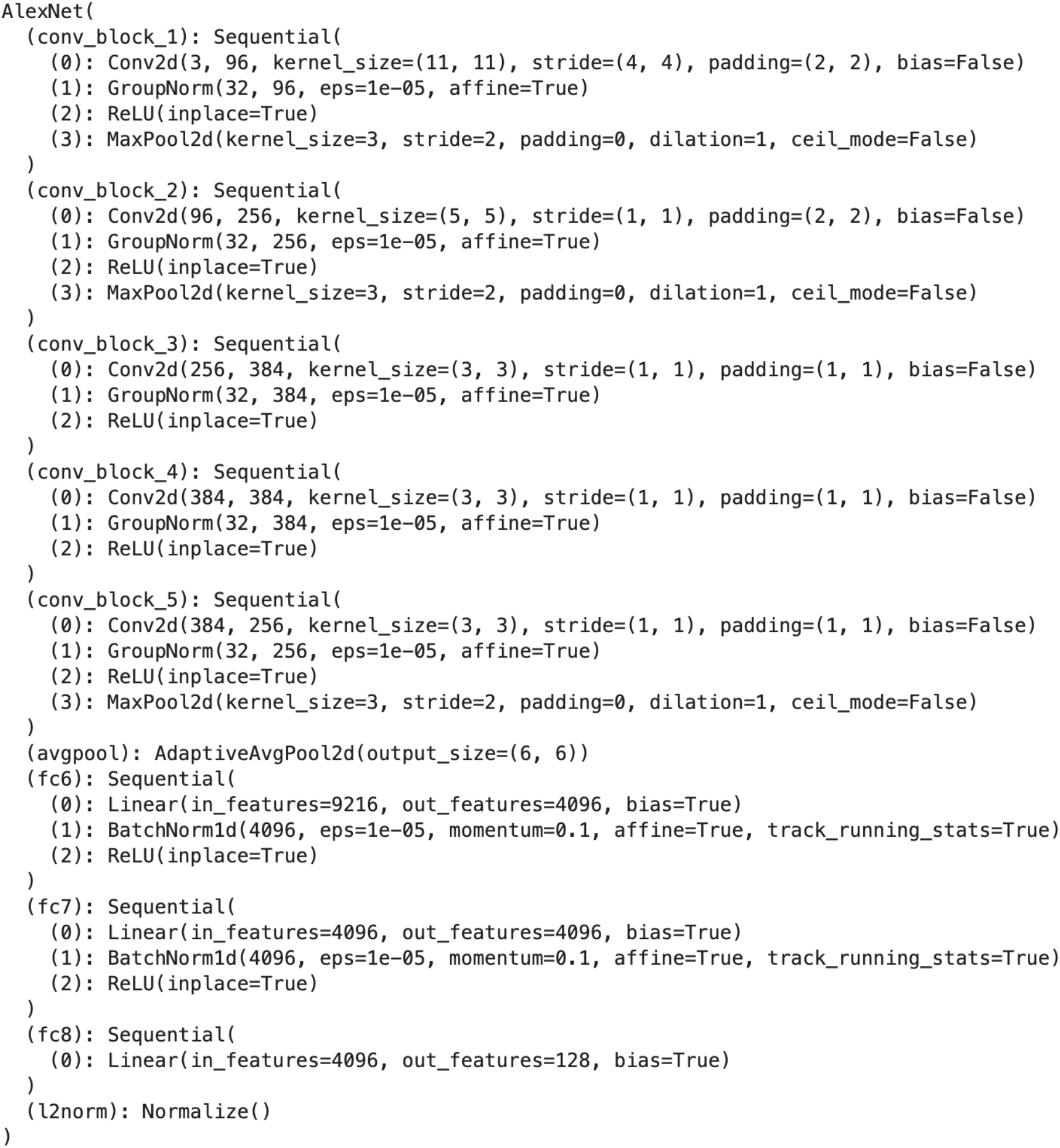
The Alexnet-gn model architecture

**Supplementary Figure 2:**
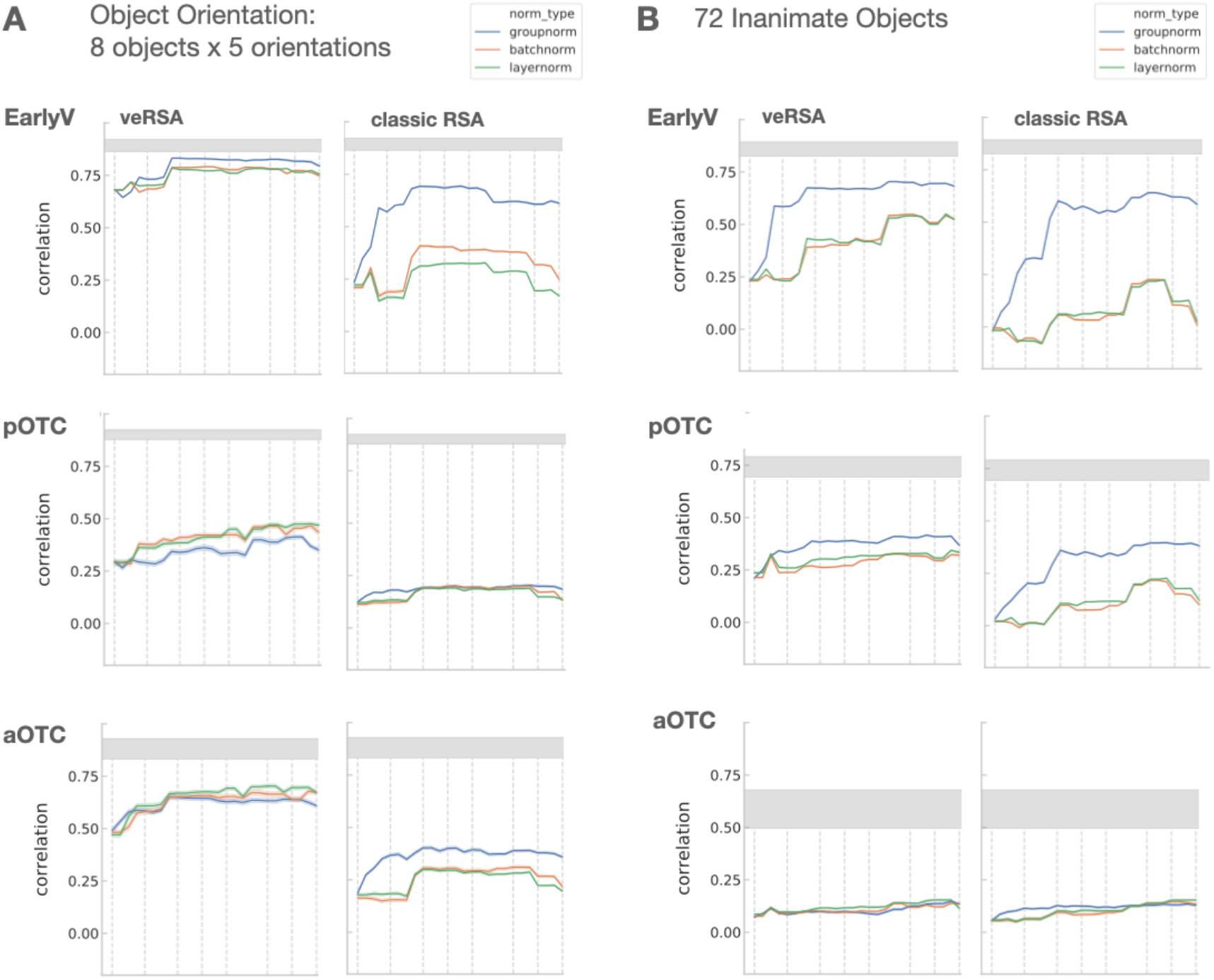
Comparison of untrained models with different normalization layers. Layerwise correlations are shown with both classic RSA and veRSA methods (y-axes), plotted as a function of model layer (x-axes), for all brain sectors (rows), and both datasets (A,B). Untrained Alexnet models with group normalization are in blue, those with batch normalization are in orange, and those with local response normalization are in green). Overall, the untrained model with group normalization layer tends to have a stronger correspondance with the brain data, compared to models with other normalization schemes.

**Supplementary Figure 3:**
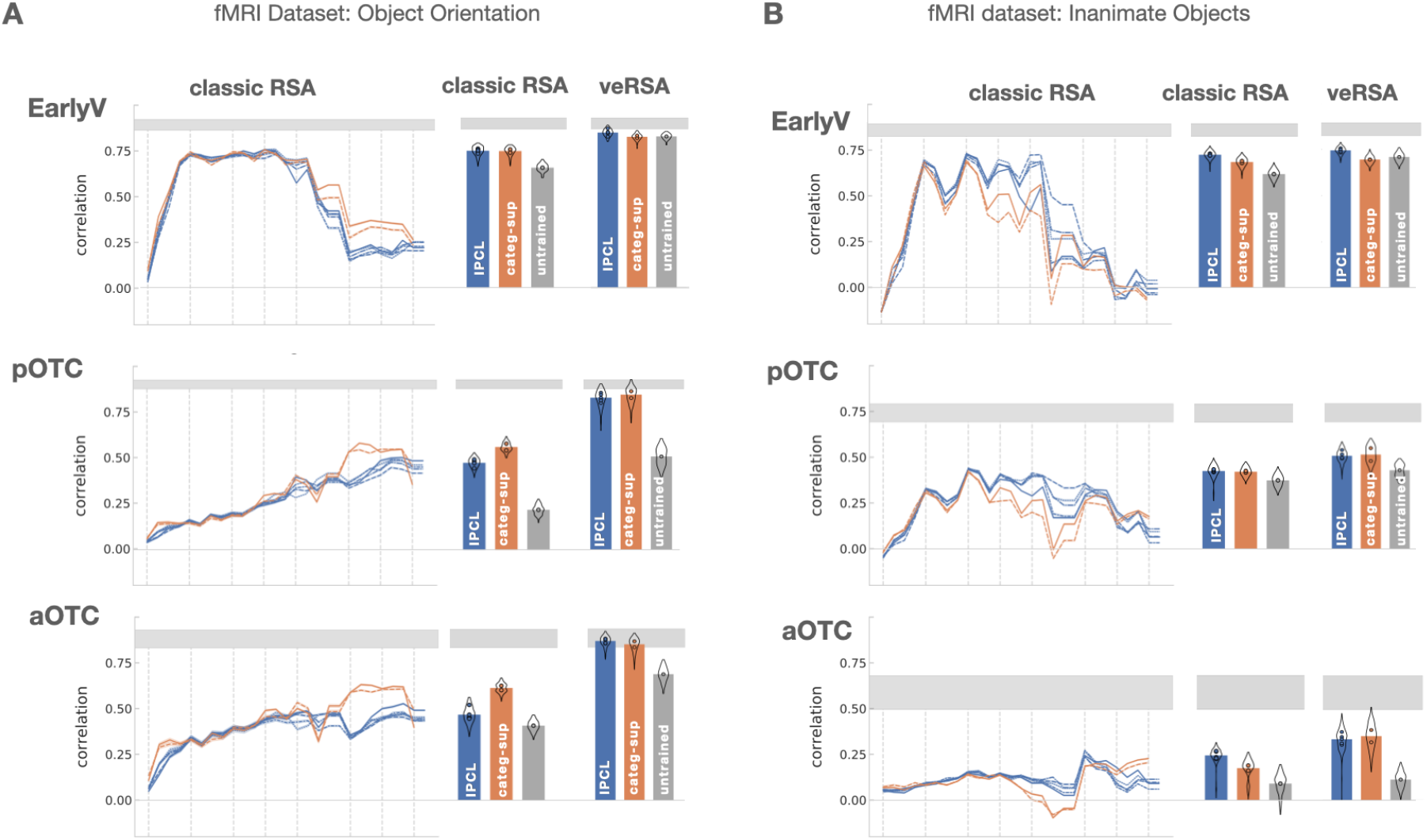
Model-Brain correspondence using Classic RSA. Layerwise correlations (y-axis) are plotted as a function of model layer (x-axis), for each brain sector (rows), in both datasets (A, B). IPCL models are in blue; Category-Supervised models in Orange. Adjacent bar-graphs plot cross-validated max correlation (y-axis) for the primary IPCL models, category-supervised models, and an untrained model. For comparison, the model-brain correlation estimated through veRSA, reported in Figure 2, is also replotted here. Overall IPCL and category-supervised models achieve similar fits to the neural data, with the exception of later model layers in pOTC and aOTC for the Object Orientation dataset. Error bars reflect a mirrored density plot (violin plot) showing the distribution of correlations across all split-halves, aggregated across instances of a given model type. Distributions are cutoff at ±1.5 IQR (interquartile range, Q3-Q1).

**Supplementary Figure 4:**
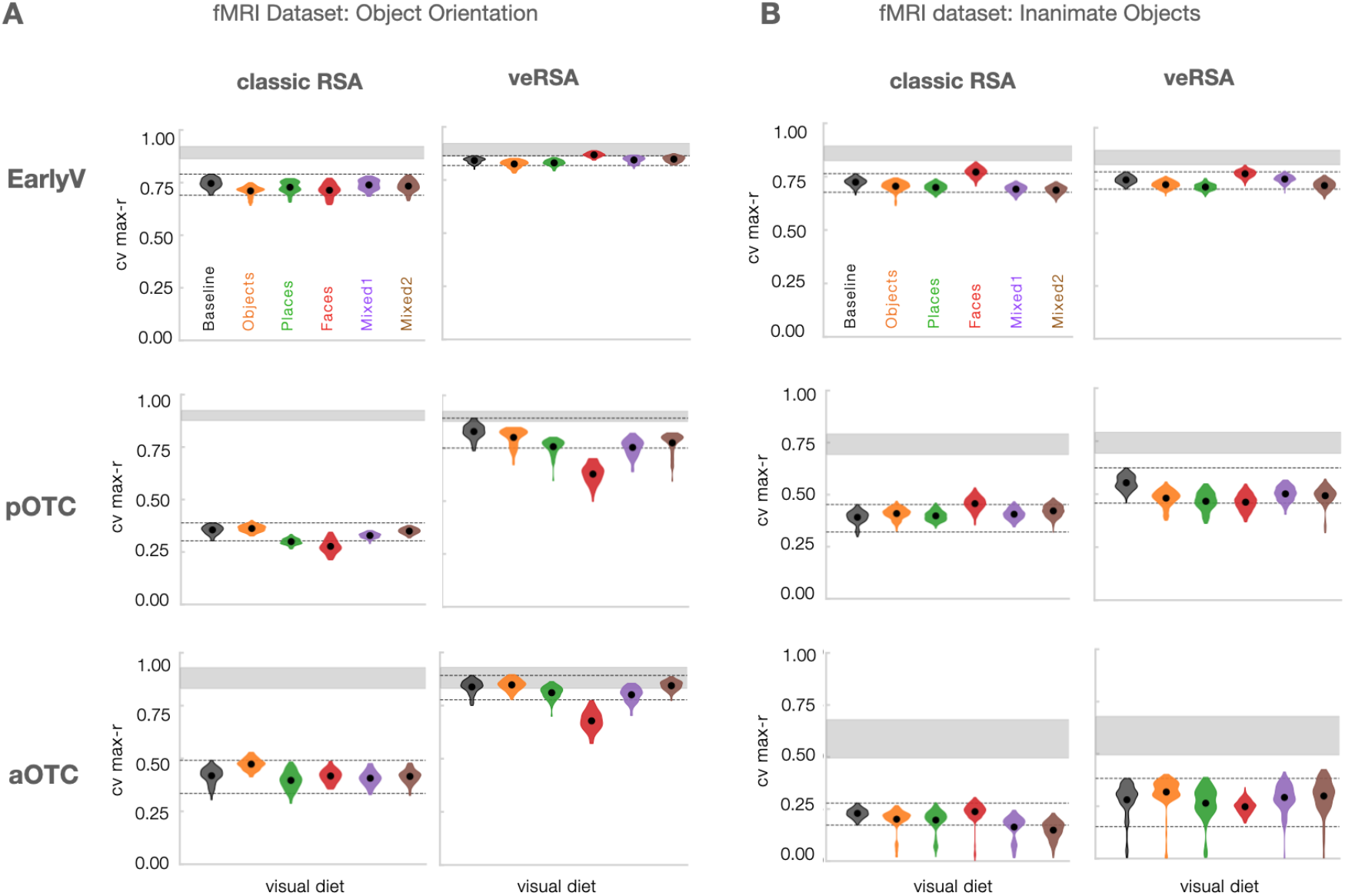
Visual Diet Variations, comparing classic RSA and veRSA. Cross-validated max-r (y-axes) was computed both with classic RSA and veRSA, in all three brain sectors (rows), and both datasets (A,B). IPCL models were trained on imagesets consisting of objects, places, faces, or mixed sets (indicated with color). Mean scores are shown with a black dot at the center of a mirrored density plot (violin plot) showing the distribution of correlations across all split-halves (distributions are cutoff at ±1.5 IQR, interquartile range, Q3-Q1). The dashed black lines indicate the ±1.5 IQR range for the matched baseline IPCL model trained on ImageNet. The correspondence between model RDMs and neural RDMs is greater for veRSA, but this benefit is muted for the face-trained models, specifically in the Object Orientation dataset.

---

**Supplementary Table 1:**
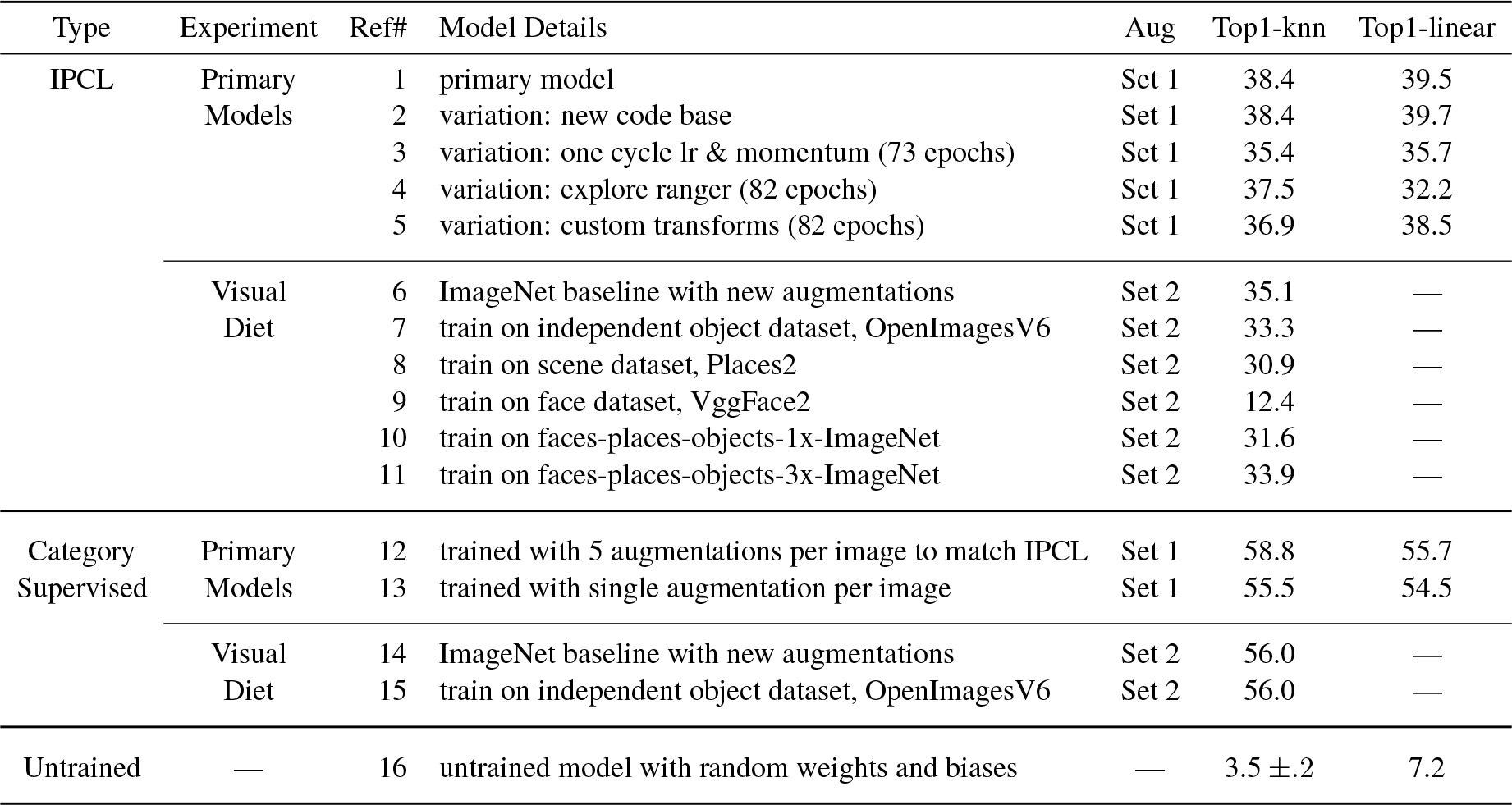
Categorization Accuracy in All Trained Models. Top1-knn classification accuracy (percent correct) is based on readout from the final layer of IPCL and untrained models (fc8), and the penultimate layer (fc7) of category-supervised models (which show higher linear readout from the penultimate layer than the final layer, as is often observed with these protocols; Chen et al., 2020a). For the untrained model, Top1-kNN shows the mean and standard deviation across 10 untrained models. Top1-linear evaluation accuracy is based on linear readout from the penultimate layer (fc7) for all models.

**Supplementary Table 2:** Statistics of Primary IPCL Models. Comparison of IPCL against the category-supervised and untrained models presented in Figure 2. Each of the five IPCL models (#1-5 in Table S1) were compared to the category-supervised models (#12,13 in Table S1), and an untrained model. Paired t-tests were computed on cross-validated max correlation values across all possible split-halves of the data (df = number of splits - 1), with a correction for non-independence of the samples (see Methods). Comparisons that were significant at the bonferroni corrected α level of .05/30=0.0017 are shown in bold. Positive t-values indicate greater predictivity for the IPCL model.

| Brain Dataset | Brain Region | Model | vs. categ-sup-12 | vs. categ-sup-13 | vs. untrained |
| --- | --- | --- | --- | --- | --- |
| Object Orientation | EarlyV | IPCL #1 | $t(69)=1.70, p=0.093$ | $t(69)=0.85, p=0.400$ | $t(69)=1.03, p=0.305$ |
| | | IPCL #2 | $t(69)=1.63, p=0.107$ | $t(69)=0.89, p=0.378$ | $t(69)=1.07, p=0.288$ |
| | | IPCL #3 | $t(69)=0.52, p=0.602$ | $t(69)=-0.40, p=0.689$ | $t(69)=-0.09, p=0.930$ |
| | | IPCL #4 | **$t(69)=3.43, p=0.001$** | $t(69)=2.69, p=0.009$ | $t(69)=2.99, p=0.004$ |
| | | IPCL #5 | $t(69)=1.90, p=0.061$ | $t(69)=1.11, p=0.271$ | $t(69)=1.36, p=0.177$ |
| | pOTC | IPCL #1 | $t(69)=-2.17, p=0.033$ | $t(69)=-0.87, p=0.385$ | **$t(69)=3.48, p=0.001$** |
| | | IPCL #2 | $t(69)=-0.34, p=0.732$ | $t(69)=1.35, p=0.180$ | **$t(69)=4.80, p<0.001$** |
| | | IPCL #3 | $t(69)=-1.24, p=0.219$ | $t(69)=0.05, p=0.957$ | **$t(69)=5.12, p<0.001$** |
| | | IPCL #4 | $t(69)=-0.56, p=0.574$ | $t(69)=0.51, p=0.614$ | **$t(69)=6.62, p<0.001$** |
| | | IPCL #5 | $t(69)=-1.89, p=0.063$ | $t(69)=-0.48, p=0.630$ | **$t(69)=5.50, p<0.001$** |
| | aOTC | IPCL #1 | $t(69)=1.39, p=0.169$ | $t(69)=0.34, p=0.733$ | **$t(69)=4.71, p<0.001$** |
| | | IPCL #2 | $t(69)=0.58, p=0.561$ | $t(69)=-0.65, p=0.517$ | **$t(69)=4.41, p<0.001$** |
| | | IPCL #3 | $t(69)=0.77, p=0.446$ | $t(69)=-0.56, p=0.577$ | **$t(69)=4.33, p<0.001$** |
| | | IPCL #4 | $t(69)=1.95, p=0.055$ | $t(69)=1.14, p=0.256$ | **$t(69)=6.13, p<0.001$** |
| | | IPCL #5 | $t(69)=1.86, p=0.068$ | $t(69)=0.97, p=0.334$ | **$t(69)=5.92, p<0.001$** |
| Inanimate Objects | EarlyV | IPCL #1 | **$t(251)=3.52, p=0.001$** | **$t(251)=3.88, p<0.001$** | **$t(251)=3.66, p<0.001$** |
| | | IPCL #2 | $t(251)=2.02, p=0.044$ | $t(251)=2.08, p=0.038$ | $t(251)=1.96, p=0.051$ |
| | | IPCL #3 | $t(251)=2.24, p=0.026$ | $t(251)=2.45, p=0.015$ | $t(251)=1.89, p=0.060$ |
| | | IPCL #4 | $t(251)=2.58, p=0.010$ | $t(251)=2.79, p=0.006$ | $t(251)=2.46, p=0.015$ |
| | | IPCL #5 | $t(251)=2.79, p=0.006$ | **$t(251)=3.20, p<0.0016$** | $t(251)=2.15, p=0.032$ |
| | pOTC | IPCL #1 | $t(251)=0.59, p=0.556$ | $t(251)=-2.37, p=0.019$ | $t(251)=2.17, p=0.031$ |
| | | IPCL #2 | $t(251)=0.45, p=0.655$ | $t(251)=-2.11, p=0.035$ | $t(251)=1.92, p=0.056$ |
| | | IPCL #3 | $t(251)=1.11, p=0.266$ | $t(251)=-1.27, p=0.204$ | $t(251)=2.47, p=0.014$ |
| | | IPCL #4 | $t(251)=2.91, p=0.004$ | $t(251)=-0.60, p=0.546$ | **$t(251)=3.93, p<0.001$** |
| | | IPCL #5 | $t(251)=0.70, p=0.483$ | $t(251)=-2.65, p=0.009$ | $t(251)=2.45, p=0.015$ |
| | aOTC | IPCL #1 | $t(251)=-0.22, p=0.826$ | $t(251)=-1.12, p=0.262$ | $t(251)=2.28, p=0.024$ |
| | | IPCL #2 | $t(251)=0.53, p=0.594$ | $t(251)=-0.91, p=0.361$ | **$t(251)=3.47, p=0.001$** |
| | | IPCL #3 | $t(251)=-0.11, p=0.915$ | $t(251)=-1.43, p=0.153$ | **$t(251)=3.30, p=0.001$** |
| | | IPCL #4 | $t(251)=1.15, p=0.252$ | $t(251)=-0.30, p=0.763$ | **$t(251)=3.59, p<0.001$** |
| | | IPCL #5 | $t(251)=0.33, p=0.741$ | $t(251)=-0.90, p=0.368$ | **$t(251)=3.22, p=0.001$** |

**Supplementary Table 3:**
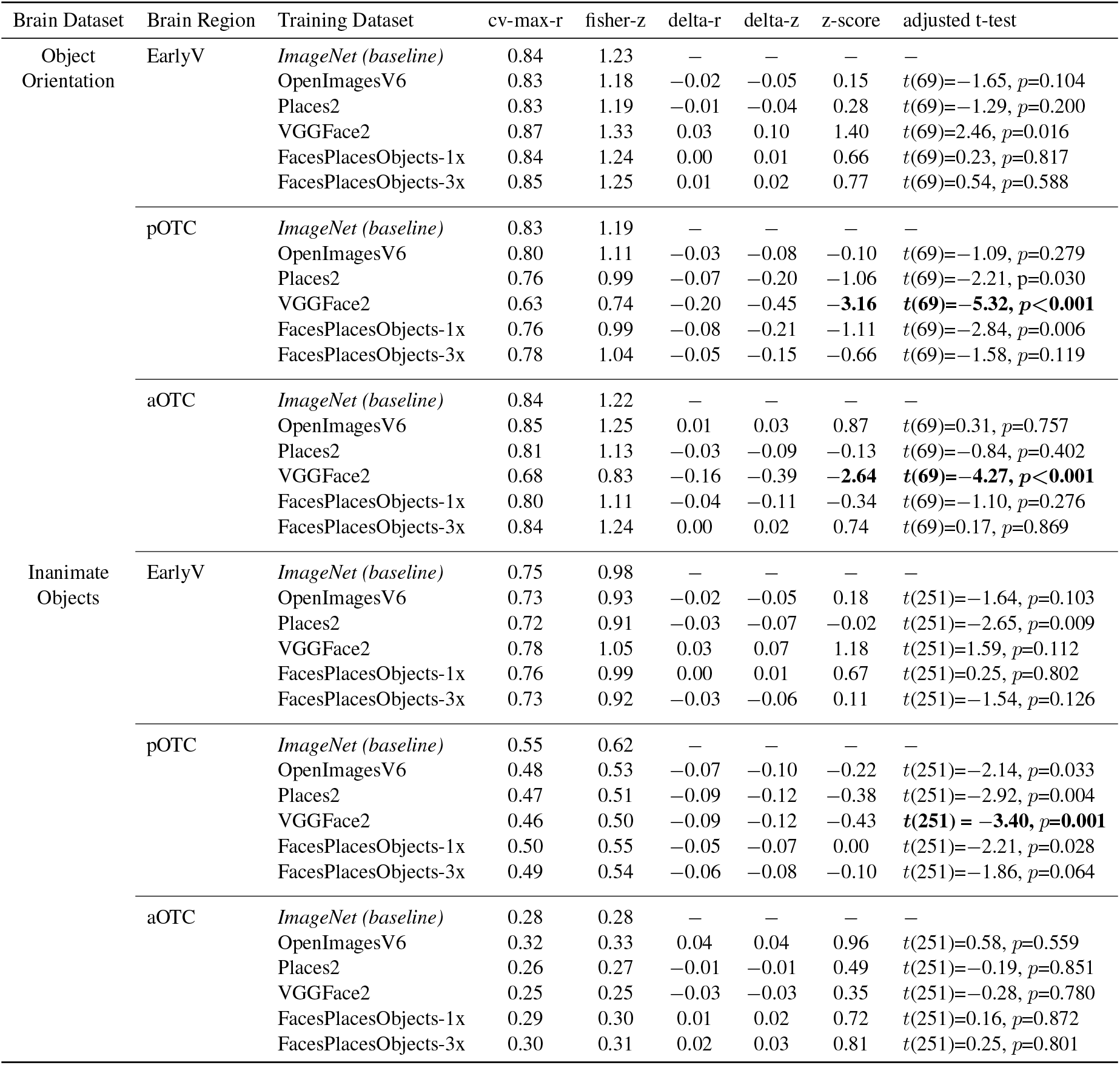
Statistics of Visual Diet Manipulation. Comparison of models trained with different visual diets against the ImageNet baseline model, presented in Figure 3. The cross-validated maximum correlation (cv-max-r) scores were fisher-z transformed for statistical analyses. The difference from baseline was computed both for cv-max-r (delta-r), as well as the fisher-z transformed values (delta-z). To quantify the magnitude of differences from the baseline, the mean and standard deviation of these delta-z values was computed and a z-score was calculated (z-score). The VGGFace2 models were outliers in their difference from baseline (z-scores >2.5 SD from the mean). We also performed paired t-tests across all split-halves of the data, comparing each model and the corresponding ImageNet baseline, and adjusting for the non-independence of the samples (see Methods). Only VGGFace2-trained networks were statistically significant different from the ImageNet baseline at the Bonferroni corrected α level of .05/30=0.0017.

## Notes

### Competing Interest Statement

The authors have declared no competing interest.

